# Traces of Meaning Itself: Encoding distributional word vectors in brain activity

**DOI:** 10.1101/603837

**Authors:** Jona Sassenhagen, Christian J. Fiebach

**Affiliations:** Department of Psychology & Brain Imaging Center, Goethe University Frankfurt, Germany

**Author notes:** Address for Correspondence: Jona Sassenhagen, Department of Psychology, Goethe University Frankfurt, PEG Theodor-W.-Adorno-Platz 6, 60323 Frankfurt, Germany. **Conflict of Interest:** Authors report no conflict of interest. **Funding Sources:** The research leading to these results has received funding from the European Community’s Seventh Framework Programme (FP7/2013) under grant agreement n° 617891 awarded to CJF.

**Keywords:** Semantics, EEG, word2vec, N400, Encoding/Decoding, MVPA

## Abstract

How is semantic information stored in the human mind and brain? Some philosophers and cognitive scientists argue for vectorial representations of concepts, where the meaning of a word is represented as its position in a high-dimensional neural state space. At the intersection of natural language processing and artificial intelligence, a class of very successful distributional word vector models has developed that can account for classic EEG findings of language, i.e., the ease vs. difficulty of integrating a word with its sentence context. However, models of semantics have to account not only for context-based word processing, but should also describe how word meaning is represented. Here, we investigate whether distributional vector representations of word meaning can model brain activity induced by words presented without context. Using EEG activity (event-related brain potentials) collected while participants in two experiments (English, German) read isolated words, we encode and decode word vectors taken from the family of prediction-based word2vec algorithms. We find that, first, the position of a word in vector space allows the prediction of the pattern of corresponding neural activity over time, in particular during a time window of 300 to 500 ms after word onset. Second, distributional models perform better than a human-created taxonomic baseline model (WordNet), and this holds for several distinct vector-based models. Third, multiple latent semantic dimensions of word meaning can be decoded from brain activity. Combined, these results suggest that empiricist, prediction-based vectorial representations of meaning are a viable candidate for the representational architecture of human semantic knowledge.

## 1. Introduction

How is the meaning of words represented in the human mind? A range of neurocognitive models discusses the neural underpinnings of semantic representation (see Borghesani & Piazza, 2018 for a recent review), focusing mostly on studying the neural localization of semantic processing (e.g., Patterson et al., 2007; Lambon-Ralph & Patterson, 2008) or its temporal organization (Lau et al., 2008; Kutas & Federmeier, 2000). Yet we have so far gained comparatively little insight into how semantic meaning is represented in our mind. From a psychological or linguistic perspective, the methods of cognitive neuroscience can however also be fruitfully used for investigating the *nature* of semantic representations. For example, functional mapping of brain activation (using methods such as functional MRI or event-related potentials/ERPs of the human electroencephalogram) has established that semantic features like the concreteness vs. abstractness of a word reflect in distinguishable neural signatures elicited during lexical processing (e.g., Fiebach & Friederici, 2004; Krause et al., 1999). Similarly, in some contexts, action-associated words have been shown to co-activate brain regions associated with the corresponding motor representations (Hauk, Johnsrude, & Pulvermüller, 2004) while concrete and imageable words co-activate sensory brain regions (e.g., Aziz-Zadeh & Damasio, 2008; Aziz-Zadeh et al., 2008). And at the level of sentence semantics, it was for example demonstrated that abstract semantic and experiential world knowledge are processed similarly (e.g., Hagoort, Hald, Bastiaansen, & Petersson, 2004).

However, rather than providing direct insights into the nature of semantic representations, these and many other neurolinguistics studies are restricted to testing indirect implications of particular aspects of theories of semantic meaning, often in specific processing contexts (e.g., during the comprehension of incongruent sentences). But theories of semantics have to account for both, the mental representation of semantic concepts and how they are processed. Our understanding of the nature of semantic representations is thus not sufficiently constrained by the mere fact that, e.g., concrete and abstract words are not processed *identically* in the human brain. Put differently, empirical work has managed to demonstrate that brains at the very least encode concreteness, but this insight does not further our understanding of how semantic information is represented. This argument extends beyond concreteness effects to all domains of semantic knowledge. Only recently have theoretical and methodological developments as well as the availability of novel datasets made it possible to address more directly outstanding questions regarding fundamental aspect of semantics, including the nature of semantic representations.

An empirical investigation of what aspects of semantic meaning are represented and how they are represented, requires that the to-be-tested models are quantitatively explicit. This is, however, typically not the case for psychological or neurolinguistic models of semantics like, e.g., classical symbolic theories of meaning (Fodor, 2004), prototype theories of meaning (Rosch, 1975), embodied cognition theories of language (Pulvermüller, 2013), or neuropsychological models like the hubs and spokes model of semantics (Lambon-Ralph & Patterson, 2008). This contrasts with the field of computerized natural language processing (e.g., Jurafsky & Martin, 2014), which has in the last years seen a very dynamic development of strongly empiricist feature-based machine learning models of semantics. Models like the so-called *word2vec* family (Mikolov et al., 2013; Pennington et al., 2013) have been successfully applied to most domains of language processing, like sentiment analysis (Felbo et al, 2017), machine translation (Mikolov et al., 2013b), or document retrieval (Ju et al., 2015). Here, we propose that machine learning techniques (Hastie, Tibshirani, & Friedman, 2009) can be leveraged to *directly* test the psychological plausibility of such quantitative theories of semantic representation by assessing their fit with neuroimaging results, and that this will advance our understanding of the nature of semantic representations.

### 1.1 Prediction-based Distributional Models and Semantic Knowledge

How do distributional models of the word2vec-family function? Mikolov, Chen, Corrado and Dean (2013) introduced a prediction-based word embedding model implemented as a simple single-layer neural network that learns to predict a withheld target word given a ten-word context (i.e., the five word strings preceding and following the target in the training corpus, independent of any grammatical constraints). This corresponds to the Continuous Bag of Words version of the word2vec model. The inverse – i.e., predicting a context from a target word – is the Skip-Gram model. As Mikolov et al. (2013) describe, both perform approximately equivalently. The input and output spaces of such networks are large: if trained on sources such as the English Wikipedia, they contain > 100.000 entries (i.e., unique words). Interestingly, their internal structure is however very simple – comprising only a few tens or hundreds of neurons (often 300; see, e.g., Fig. 1A for a schematic illustration of the model architecture). We hypothesized that this ‘many-to-few’ compression may resemble the manner in which the human brain learns and represents semantic knowledge. Accordingly, we here test the relationship between brain signals elicited during word processing and the vectorial representations of these words resulting from empiricist learning models like word2vec.

**Figure 1:**
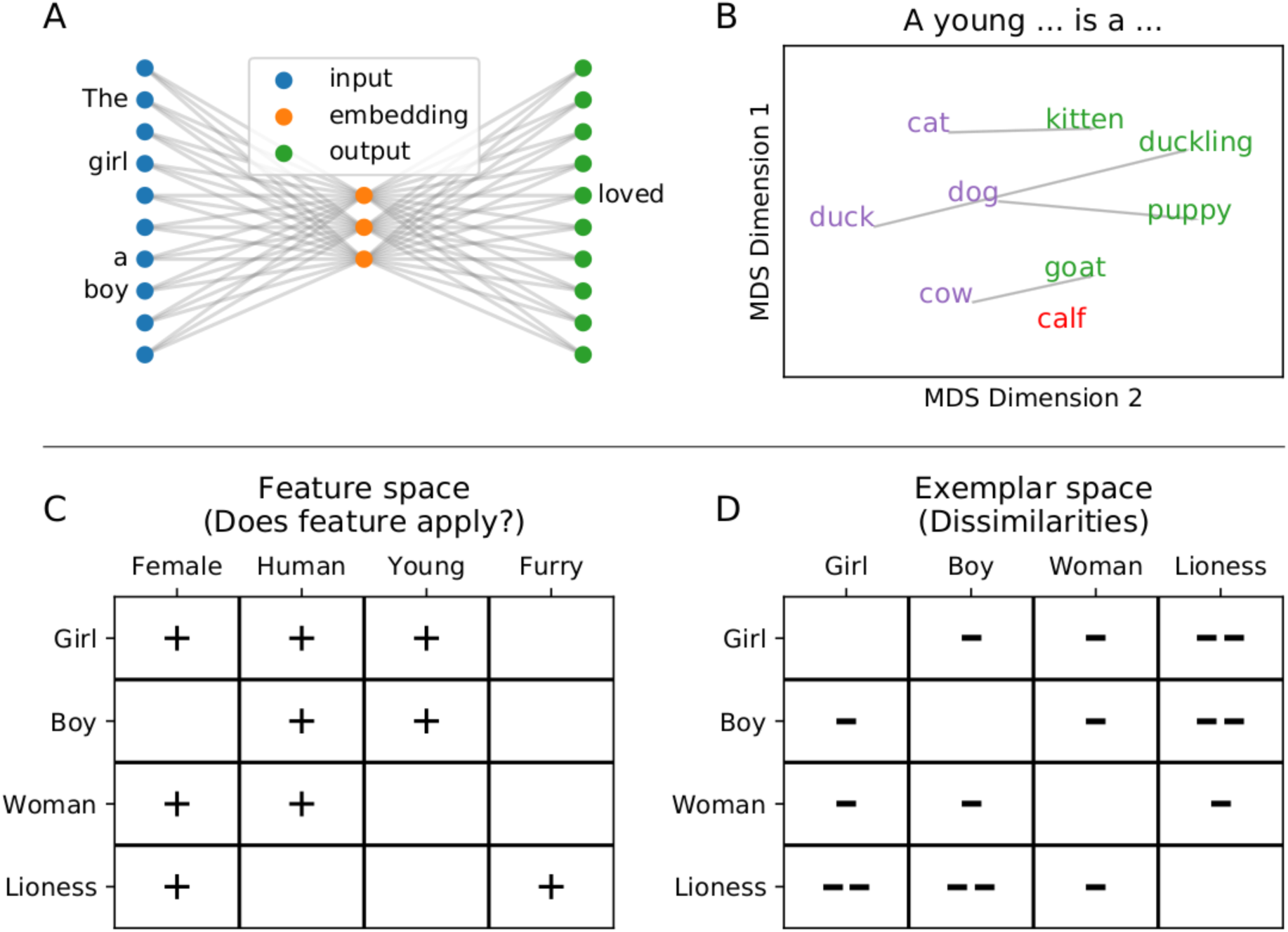
Vectorial models of semantic word knowledge. Top: Architecture and example content of distributional models. A: Basic architecture of a so-called Continuous Bag of Words prediction model for word embeddings. An input layer (blue) receives a context, and aims to learn weights on the internal layer (orange), which in turn allow the prediction of the omitted target word (green); read from right to left, this corresponds to a Skip-Gram Model. B: Vectorial models can learn semantics-like relationships. For purposes of illustration, we looked up vectors V in a pre-trained vector model (Mikolov, Grave, Bojanowski, Puhrsch, & Joulin, 2017), for four animals (*cat, dog, cow, duck).* We then added to them the vector for the word *young*, and retrieved the word closest to the resulting vector, resulting in *kitten, puppy, goat,* and *duckling*, respectively. Only for the word cow, this calculation of V_cow_ + V_young_ did not result in a young instance of the species cow (i.e., a calf). The model, accordingly, is capable of roughly representing semantic relationships such as *‘a young W_1_ is a W_2_’*. For visualization purposes, these eight words plus the missing response ‘*calf’* were embedded into two dimensions with Multidimensional Scaling as implemented in scikit-learn (Pedregosa et al., 2011). C,D: Demonstration of feature (C) and exemplar-based (D) representations of concepts. “+” indicates the presence of a feature (left). The number of “-”s represents the dissimilarity between two concepts (right).

Artificial learning systems, under which distributional models of semantics can be subsumed, resemble human learners in that in the interest of generating abstract, generalizable inferences, they benefit from constraints on their capacity (Elman, 1991). This is also a crucial factor for word2vec-family models, which perform better when the internal network size is smaller (Landauer & Dumais, 1997; Mikolov et al., 2013). Under such situations of limited internal ‘representational capacity’, the prediction-based training regime forces the internal layer, which is unable to simply ‘memorize’ all mappings, to discover latent semantic dimensions (Landauer & Dumais 1997; Baroni et al., 2014). This ‘compression’ is essential to word embedding models. Consider, as an example, the relationship between the (African) lion and the (Asiatic) tiger, who rarely appear at physically identical places in their natural environments. An unconstrained representation (i.e., a manually annotated expert system like the WordNet of Miller, 1995) would represent isolated facts like “lions are mammals” (or +MAMMAL), “tigers are mammals”, “lions are quadrupeds”, etc. (see Fig. 1C for schematic). In contrast, a prediction-based and capacity-constrained distributional model would have to identify what is shared between lions and tigers (e.g., that they are large cats, occur in similar text contexts, etc.) and code this information in an abbreviated, synergistic fashion. This strategy of reducing a dataset, such as the co-occurrence statistics of words, to a limited number of latent, generalizable factors underlying its axes of variance, is particularly well suited for uncovering such implicit, underlying similarities. These *distributed and distributional* models are indeed capable of learning word meaning from distributional information only (see also Landauer & Dumais, 1997). As Fig. 1B schematically demonstrates, they are even capable of tasks such as compositionality (see also Mikolov et al., 2013), which is generally considered a key aspects of any theory of semantics (Fodor & Lepore, 1999).

### 1.2 Vectors in the brain

After training a distributional network model, the weights between each word and the – e.g., 300 – neurons in the hidden layer are understood as an *embedding* of the word, so that each word is described as a vector of 300 values (neural weights) corresponding to the axes of the vector space. This is equivalent to a position in a high-dimensional state space (Churchland, 1993). The vectors are dense and real-valued. That is, unlike typical hand-designed feature space models in which only very few words carry a feature such as, e.g., +HUMAN or +QUADRUPED, in a typical word embedding every word ranks somewhere on every axis. These axes are generally not directly interpretable, and do not typically correspond to natural-language concepts (like animacy). Instead, semantic knowledge is coded in a distributed manner, so that many axes will each be partially correlated with any given semantic dimension (Landauer & Dumais, 1997). One way for making visible the semantic knowledge in these models is to project them into exemplar space, e.g., by computing the correlation between all items. The resulting item-by-item representation shows how similar each concept is to all other concepts (see 1D), which corresponds to a *kernel*-based representation of the semantic space (see below).

Embedded distributional models are highly effective in accounting for human processing of words in contexts. More specifically, they can predict priming effects (Günther et al., 2016a, 2016b), human behavioral performance in multiple other psycholinguistic tasks (Mandera, Keuleers, & Brysbaert, 2017), semantic association ratings (Rubinstein, Levi, Schwartz, & Rappoport, 2015), similarity ratings (Mandera et al., 2017), and ERP measures of the fit of a word with its context (Broderick, Anderson, Di Liberto, Crosse, & Lalor, 2018 Ettinger, Feldman, Resnik, & Phillips, 2016). But that is almost by design, as distributional models are trained by finding patterns in item-context pairs (Levy & Goldberg, 2014), and the observation above, i.e., that context effects provide only indirect insight into the nature of semantic representations, also applies to empirical tests of distributional models of word semantics as long as they are based on item-context associations. In the present study, we go one step further and explore whether distributional models can directly account also for the neurocognitive *representation* of word meaning in the human brain. To this end, we postulate that understanding the meaning of a word is equivalent to transitioning the brain into a (more or less short-lived) unique state that systematically depends on the meaning of the perceived word. If this hypothesis is true, it should be possible to: First, predict not only behavior (see above), but also brain activity based on word vectors; second, predict not only item-context or item-item effects, but also neural correlates of context-free word processing; and third, conversely, partially recover a word’s position in vector space from brain activity elicited by the respective word.

### 1.3 Relation to prior work

A number of recent studies have used vectorial models to identify neural correlates of sets of semantic categories or features (e.g., Kemmerer, Castillo, Talavage, Patterson, & Wiley, 2008; Mitchell et al., 2008; Sudre et al., 2012; Xu, Murphy, & Fyshe, 2016; Pereira et al., 2018; Gauthier & Ivanova 2018; and Wehbe et al., 2014). Recently, Huth, de Heer, Griffiths, Theunissen, and Gallant (2016) have for example shown that it is possible to predict with high accuracy brain activity elicited during listening to narratives, by using as predictors the position of the corresponding words in an implicit state space based on taxonomic labels (Miller, 1995). This work has revealed a distributed ‘tiling’ of semantic features across the cortex, and a particular sensitivity to categories such as the social relevance of words.

In contrast to this seminal work, we here aim at using the framework of distributional models to examine the neural representation of word meanings *themselves* – i.e., the neural signatures of words bereft of context, and the hypothesized systematic relationship to their position in semantic vector space as quantified with distributional models learned in an empiricist fashion from statistical patterns of co-occurrence alone. Specifically, we tested whether word-associated brain activity can be predicted – *encoded* – from word2vec-style distributional vector representations of these words. We found that different vector spaces could successfully be encoded in brain activity. Secondly, we explored as a proof of principle what information about word vectors is contained in the electrophysiological activity of the brain elicited during word processing, by testing whether the word’s position in (for the sake of interpretability, dimensionality-reduced) vector space can be *decoded* from brain activity. Then, we established conceptual labels for the aspects of word vectors decodable from brain activity. This latter stage was conducted to ensure that brain activity was correlated with the *semantic* information in distributional models, not (or at least not exclusively) with other information about words that these models might contain (e.g., about word frequencies or similar lexical properties of words). We indeed observed that for components corresponding to semantic dimensions, vector-space factor scores of words could be read out from brain activity. Taken together, the work presented here is an initial demonstration that it is plausible that brain states elicited by words are approximately isomorphic to the position of these words in a distributional model of word meaning.

## 2. Methods

### 2.1 Datasets and Preprocessing

We here report results from two datasets, acquired in two different languages and using two different tasks: The first dataset contains ERP data for 960 visually presented English nouns (n=75; 28 EEG channels; for details see Dufau, Grainger, Midgley, & Holcomb, 2015). Words were shown on a screen for 400 msec, followed by a 600 msec blank screen, and were mixed with 140 nonword probes requiring a manual response (discarded from analysis). The second dataset was acquired in our own lab and involves EEG signals elicited by 150 visually presented German nouns (presented for 1,000 msec following a 500 msec fixation cross; n=35; for details see [a link to a preprint for a manuscript which we here withhold for the sake of doubly-blind peer review]). In this experiment, for each word, participants were instructed to press a button if and only if it was synonymous with the previous word (the 10 synonymous probes requiring a button press were ignored in this analysis). EEG was collected from 64 ActiCap active electrodes via a Brainamp amplifier (Brainproducts; Gilching, Germany).

Data analysis was conducted in MNE-Python (Gramfort et al., 2013). The English dataset was obtained from the original authors in a preprocessed form, in the form of per-word average ERPs across subjects (-100 to 920 msec post word onset; see Dufau et al., 2015). For the German dataset, eye movement artefacts were removed via ICA (Jung et al., 2000), and a .1-30 Hz bandpass filter was applied. Both datasets were downsampled to 200 Hz. Then, analogous to the English dataset, average ERPs across subjects were calculated individually for each word for the German data. Compared to single-subject analysis and averaging, this approach leads to higher signal-to-noise ratios, but does not change the overall pattern of results.

We note informally that we have applied the same analysis to a series of other datasets; the results we show here can be replicated on a number of other sufficiently large datasets of ERPs elicited during single-word presentation.

### 2.2 Word Embeddings

For all following analyses, for the vector semantics, we relied on FastText, i.e., a further development of the above-described word2vec algorithm (Bojanowski, Grave, Joulin, & Mikolov, 2016). We used the implementation from the natural language processing package GenSim (Řehŭřek & Sojka, 2011) and for both German and English, the publicly available, pre-trained, 300-dimensional embeddings provided by Mikolov et al. (2017). Note that FastText’s primary improvement upon word2vec is that by accounting for units below the word level, it naturally handles inflected forms. Due to this, it can better account for morphology-rich languages like German. Further machine learning (de-/encoding) was conducted via sklearn (Pedregosa et al., 2011) and visualization done with Seaborn (Waskom et al., 2018), both using the Python programming language.

### 2.3 Encoding Semantic Vectors in Brain Activity

We attempted to model the dependency of brain states on the position of the respective word in a semantic state space via an encoding model (King et al., 2019). Specifically, we predicted brain activity by assuming that the neural state after having processed one word resembles a sum of neural states after having processed other words, weighted by the similarity of these words to the target word (if viewed as a ’dual‘ problem; see below), or as a linear weighted sum of the features (vector-space axes). To provide a baseline model, we also conducted the same analysis based on a non-distributional model of word meaning. For this, we chose WordNet (Miller, 1995), a well-established database of lexical/semantic relations which contains a manually constructed taxonomy of words (also used, for example, in recent neurocognitive work by Huth et al, 2016).

For both languages, within a 10-fold cross-validation loop, a Ridge regression (i.e., a linear regression with *L*_2_ norm regularization) was trained to predict the entire pattern of brain activity elicited by a word, across all measurement channels, using the respective word vectors as predictors. On each fold, the regression model used 90% of the dataset as training trials to learn one coefficient for each of the vector-space model’s 300 dimensions, for each combination of time point and sensor. Subsequently, EEG activity (i.e., the pattern of signal amplitudes across sensors) was predicted for words not seen during training (i.e., the 10% trials held-out in the respective fold), based on the dot product of the regression coefficients and the vector-embedding of the respective word. To account for the high temporal resolution of EEG, in each fold, independent regression models were fitted for each time point, and predictions on the test trials were also made per time point. The quality of these predictions were scored using the signed, squared correlation between predicted and observed EEG activity (i.e., ERPs), which provided an estimate of the systematic isomorphism between word vectors and electrophysiological brain activity.

These correlation coefficients were concatenated, resulting in one time series (from -100 to +920 msec time-locked to stimulus onset) per fold corresponding to how well brain activity at each time point in the trial can be explained as a linear weighting of the word vectors for the respective word. These time courses were then averaged across folds, and 95% bootstrapped confidence intervals across folds were calculated for each time point. For a visualization of this procedure, see Fig. 2. For statistical evaluation, first, the resulting prediction accuracy values were averaged in the N400 time window (300-500 msec; Lau et al., 2008), and a Wilcoxon signed-rank test against chance was conducted across folds. To control for potential biases, this was repeated for the average of the entire data epoch.

**Figure 2:**
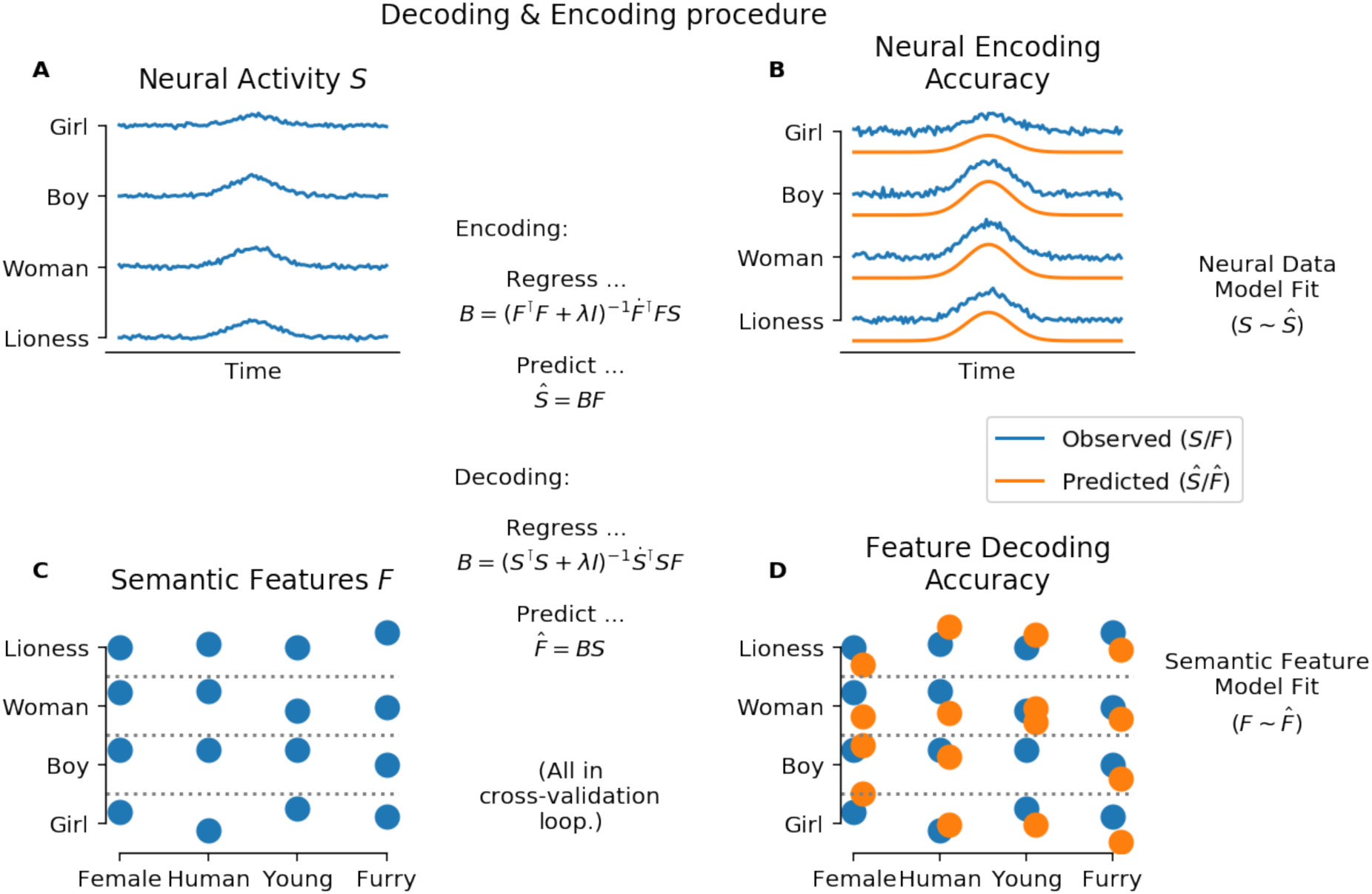
De- and Encoding procedure, didactic example for simulated values. Top: Encoding procedure. (A) Brain activity source data. For each word, the y axis represents neural activity (i.e., microvolts in the case of EEG (ERP measurements). (B) Comparison of predicted (orange) to the observed (blue) data. Center: A predictive model B is fit by estimating the coefficients minimizing a least-squares equation, to predict the ERP pattern (S) from the words’ semantic vector embedding F. Bottom: Decoding procedure. (C) Interpretable semantic feature rankings of word vectors. Here, the y axis represents the ranking on each of four exemplary semantic dimensions. (D) Actual (blue) are compared to predicted (orange) semantic features, for each word. Center: In the decoding analysis, a predictive model B is fit by estimating the coefficients minimizing a least-squares equation, to predict semantic features F from ERP signal S.

As noted, to compare the distributional word embedding model against a reasonable baseline model, we repeated the same analysis with similarities based on the WordNet taxonomy (Miller, 1995) for the English dataset and, for the German dataset, using a German equivalent (GermaNet; Hamp & Feldweg, 1997). WordNet’s taxonomy corresponds to hyponymy/hyperonymy relations – i.e., “is-a” relationships like ‘a robin is a bird’. To predict EEG/ERP activity from this taxonomic model, we calculated *path similarities* for each pair of words, i.e., the distance to the next shared hyperonym or hyponym. This lead to a 960 × 960 matrix (English) and a 150 × 150 matrix (German), respectively, of semantic word/word distances which was then used to predict EEG/ERP patterns across time points and electrodes and for checking the fit with actual ERPs on unseen trials, analogous to the FastText-based encoding described above (i.e., 10-fold cross-validation).

We calculated the difference, within each language, between prediction accuracy scores for the distributional/FastText-based predictions minus the score for WordNet- or GermaNet-based predictions. This was calculated separately for each fold, and a Wilcoxon Rank-Sum test on the average of the differences within each fold was conducted to establish which model yielded better predictions of the ERP patterns: the explicit taxonomic (i.e., WordNet/GermaNet) model, or the distributional model. Because the feature space built from Wordnet is initially larger than that for FastText vectors for English (960 items × 960 items vs. 960 items × 300 dimensions) and smaller for German (150 × 150 vs. 150 × 300), to not bias the procedure in any direction, we computed this contrast also on a transformation of the FastText vector predictor array (*XX*^⊺^), which has the same item × item dimensionality as the Wordnet-derived matrix of path similarities. This yielded the same results (for possible reasons, see below the discussion of the Representer Theorem).

Next, to rule out that successful encoding is the result of lexical but not than semantic features being reflected in brain activity, we estimated if some of the encoding benefit for the superior model might reflect well-established psycholinguistic features like the frequency of occurrence of a word in a language, rather than the hypothesized greater similarity to the underlying neurocognitive correlates of semantics. To explore this alternative, we repeated the described analyses, but included in an exemplary manner word frequency (log counts per million as given by Dufau et al., 2015, for the English data; SUBTLEX log frequency, Brysbaert et al., 2011, for German) and concreteness (imaginability ratings from Dufau et al., 2015, for English and from Vo et al., 2009, for German) to the predictor arrays, and investigated if this influenced any of the observed effects. By doing so, we tested if any contrast between FastText-based and WordNet-based encoding performances was diminished as a result of adding explicit word frequency and concreteness values as predictors. We did not directly compare the predictive power of lexical variables like word frequency or concreteness to vector representations because this would entail comparing a very large predictor matrix (e.g., 960 predictors in English) to a much smaller one (i.e., with 2 predictor variables), so that much of the resulting differences might reflect the capacity of the algorithm to handle large feature spaces. In contrast, adding these psycholinguistic predictors to the set of baseline and word vector predictors and evaluating performance differences at the level of alternative models, reflects a meaningful test of the value of the word vector model.

Finally, to to assess how specific our results were for our specific choice of vector space, we repeated the encoding analysis for a set of other commonly used, pre-trained distributional vector spaces, based on different algorithms (i.e., not from the word2vec family), different training corpora, and different vector sizes. Specifically, we encoded the original Wikipedia-based word2vec vectors released by Mikolov et al. (2013), the movie subtitle-based vectors by Mandera et al. (2017; Subtlex), and the full set of English vectors from Pennington et al. (2013) known as GloVe, including web and Twitter corpora. These embeddings encompassed multiple vector lengths (25-300).

### 2.4 Decoding

In addition to encoding, a correspondence between the structure of brain activity and semantic spaces can also be established by decoding – i.e., by reading out properties of the stimulus based on brain activity. In our case, we tested if we could predict a word’s position in word2vec space based on brain activity (see Fig.2, bottom).

As the signal to noise ratio of both the ERPs and the word vectors is low, predicting the absolute position of a word in high-dimensional vector space would lead to low fits and would accordingly be hard to interpret. We therefore first conducted dimensionality reduction on word vectors. For this, the vector space was transformed via Kernel PCA with a cosine kernel (cosine similarity is the preferred distance measure for vector spaces, cf. Mikolov et al. 2013; see below for a discussion of kernels in vector semantics). We arbitrarily decided to investigate the first eight components (as all further components explain less variance and are more noisy and harder to interpret or read out of brain activity). I.e., to investigate to which degree brain activity contains information about word semantics, we predicted each word’s loading on eight factors, resulting from Kernel PCA reduction of word vectors, from brain activity.

More specifically, we attempted to read out from brain activity elicited by words presented in isolation (same data as used for encoding analysis) the position of each word on each of these eight factors, using as predictors ERP time courses between 300 and 500 msec (reflecting the typical N400 time window; see above). In a 10-fold cross-validation loop, Ridge regression was used to predict the position of each word in Kernel PCA-compressed vector space. Scoring was again accomplished via the signed, squared correlation (across items) between predicted and actually observed outcomes, resulting in 10 × 8 *r*^2^ values – one per fold and factor – for each language (i.e., for each of the two datasets). For each factor, a 99.375% bootstrapped confidence interval was calculated – corresponding to a Bonferroni correction for the eight factors. In addition, a time-resolved multivariate decoding analysis (King & Dehaene, 2013) was conducted by repeating the same procedure as above, but now separately for each sample time point of the full EEG data (with only electrode as feature). This resulted in 10 × 8 *r*^2^ values for each time point, for each language. 68% bootstrapped confidence intervals were calculated (not corrected) for visualization only.

### 2.5 Interpreting Word Vector Scores

As noted, the axes of distributional vector spaces do not a priori have interpretable labels, and are in fact hard or impossible to label. Doubtlessly, the vector space as a whole contains a lot of semantic information about conceptual characteristics (like +ANIMACY). But this information is latent and must be actively inferred (which is however often not easily possible). In addition, semantic information is encoded in a distributed fashion; any semantic feature, such as animacy or concreteness, correlates with multiple axes of the vector space, but the specific axes themselves are not in a one to one relationship with any semantic trait. Thus, the decoding procedure described above could not – regardless of the success of decoding –indicate that information about word *meaning* is contained in brain activity. Any factor score which could be read out from brain activity could correspond to a number of non-semantic aspects of word meaning also recoverable from vector spaces.

To rule out that a success in decoding vector-space position from brain activity may depend upon nonsemantic aspects of word knowledge, we set out to label the eight factors used in the decoding analysis. Three trained linguists considered the seven words scoring most positively and most negatively on each of the eight components, separately for each language. Each linguist proposed one interpretation; then, each linguist ranked all three proposals for each axis. The proposal with the highest mean ranking was selected as a tentative label for the factor. Note that the specific labels are not important per se; the label interpretations are subjective in nature, and no transformation of the data is guaranteed to reproduce the ‘true’ latent factor structure. Instead, this labelling was done simply to establish if brain activity allows decoding of *semantic* information at all – or if the correspondence between brain activity and distributional word vectors is due to nonsemantic information contained in word vectors. That is, if brain activity allowed reading out the position of words on factors corresponding to conceptual distinctions, then this would speak for the decoding procedure succeeding because semantic information is encoded in brain activity.

### 2.6 Visualizing the spatial structure of encoding and decoding

Lastly, we set out to exploratorily visualize at which electrodes the signals on which encoder and decoder models operated were most clear. For this, we focused again on a time window of 300 to 500 msec post stimulus onset. We excluded the English dataset as it was recorded with very sparse electrode coverage, and focused instead on the high-density German data. First, we aggregated the encoding performance over time within each electrode, and visualized the result as an interpolated scalp map. Conceptually, this corresponds to a map of where the encoder procedure works best, i.e., where whatever it is encoding is most directly represented. Next, we retrieved normalized regression coefficients for each Kernel PCA component from the decoding analysis, and similarly visualized them as scalp maps. Conceptually, this corresponds to sites where higher activity predicts a higher (or lower, depending on the sign of the coefficient) component score, i.e., where EEG/ERP activity is predictive of the respective component’s score per word.

## 3. Results

### 3.1 Encoding Semantics in Brain Activity

All four models (for English: FastText and WordNet path similarities; for German: German FastText and GermaNet path similarities) succeeded in predicting EEG activity, in particular around the N400 time window which is known to reflect semantic memory access (see Fig. 3, left; approx. 250 msec to 550 msec). The vector-space models predicted activity well in our a-priori predefined time window for statistical analyses (i.e., mean of 300-500 msec: English, *r*^2^= 0.05, SD = .025; German, *r*^2^= 0.07, SD = 0.045; *both p* < 0.005), but also when aggregating over the entire ERP epoch (both *p* < 0.005). Importantly, vector-space models predicted brain activity significantly better than the taxonomic (WordNet) models (*p* < 0.01; compare dashed vs. solid lines in Fig. 3). The right panel in Fig. 3 displays the time-resolved improvement of semantic encoding based on vector space as compared to taxonomic models.

**Figure 3:**
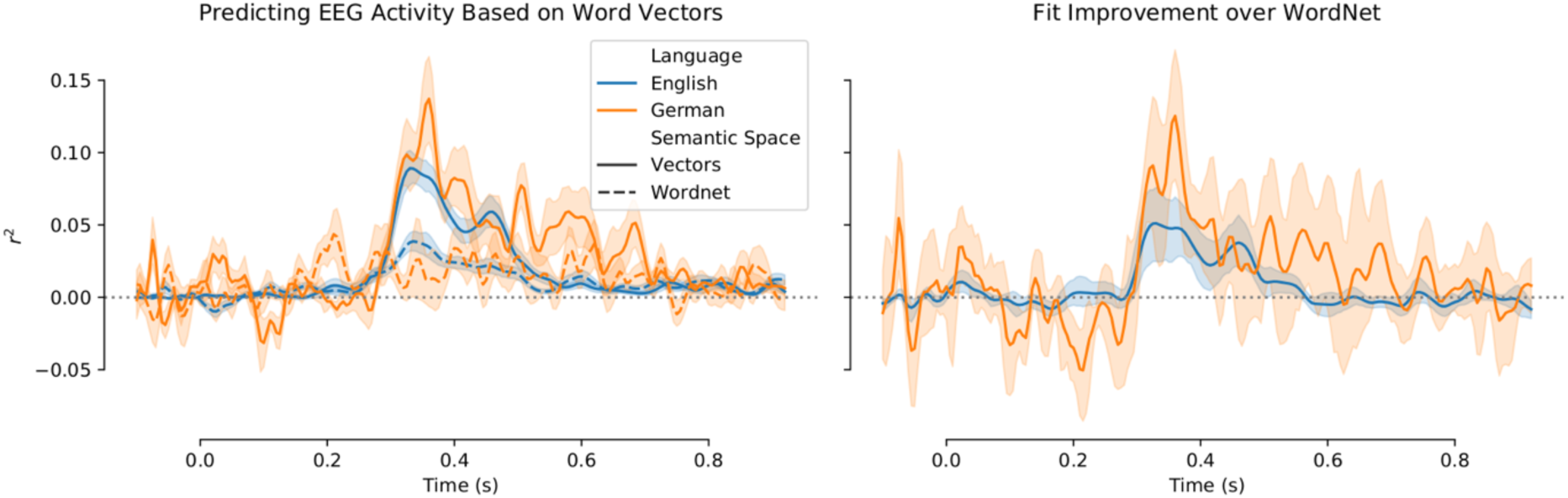
Encoding model fit for both languages. (A) Goodness-of-fit across time, displayed as the squared and signed correlation between predicted and observed EEG signals, aggregating across sensors, participants, and folds, for word vector embedding models (solid lines) and the WordNet (i.e., control) model (dashed lines) in English (blue) and the equivalent German models (orange). (B) The difference between word vector models and taxonomic models demonstrates the significantly better prediction of EEG/ERP activity from word vector models. Lines represent the mean (squared, signed) correlation between the predicted and the actual ERP. Shaded areas reflect 68% bootstrapped confidence intervals over folds.

Results for the alternative vector spaces are qualitatively similar (see Fig. *4*), indicating that distributional vectors in general allow prediction of brain activity. Interestingly, shorter vectors appeared to allow superior encoding, potentially indicating overfitting for high-dimensional vectors.

**Figure 4:**
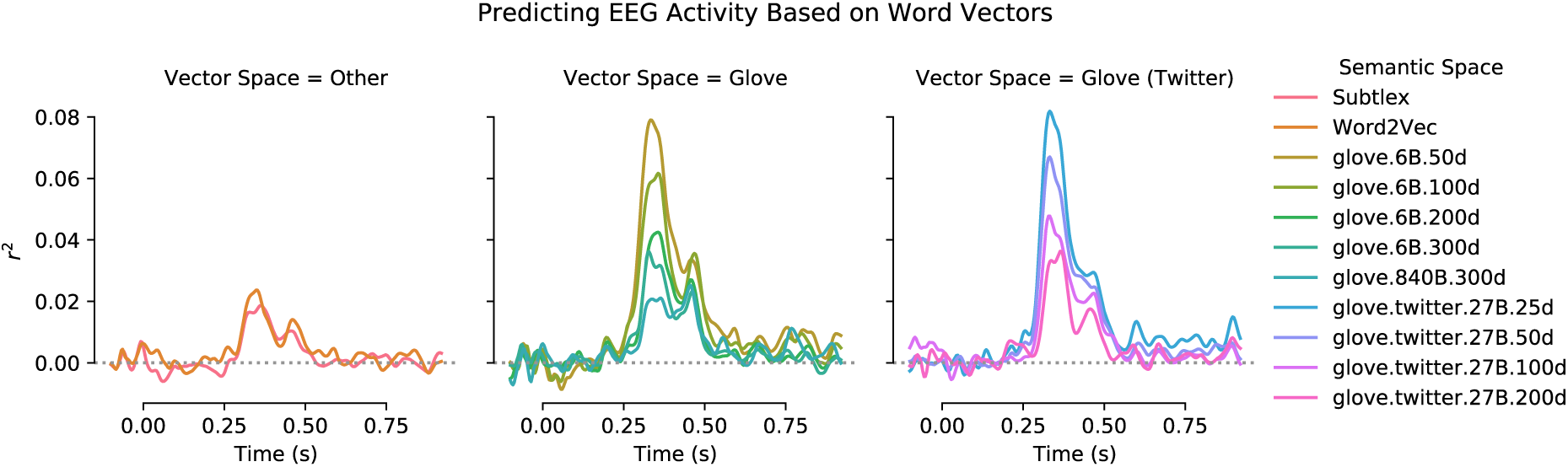
Encoding model fits for alternative (English) word vectors. All details as in Fig. 3, but each plot shows the time series for encoding quality for multiple vector space models (see right legend). word2vec- and Subtlex-vectors reflect an older algorithm and a different training corpus (movie subtitles), respectively. GloVe vectors (of various lengths) correspond to a related, but distinct algorithm (Pennington et al., 2013). GloVe vectors trained on twitter data reflects yet another corpus. Note that while encoding accuracies vary across models, all models allow above-chance prediction of activity in the N400 time window.

### 3.2 Decoding

Observing that encoding – i.e., the prediction of brain activity during word reading based on the position of these words in vector space – is possible, we set out to more thoroughly test if this was due to the semantic content of word vectors. For this, we conducted a factor analysis (Kernel PCA) of word vectors (to support decoding and to render the results more interpretable), and then attempted to decode a word’s position in (factorized) vector space – i.e., its loading on the components – based on brain activity. We also, subsequently, aimed at inferring what these factors correspond to conceptually (see next section for details).

Decoding was used to predict the ‘loading’ of each word on each of these eight factors, based on EEG activity. Results indicate that loadings on multiple factors could be read out from brain activity, with cross-validated (over 10 folds; see Methods) *r*^2^ scores between predicted and observed factor scores > .1 (*p* < .05, Bonferroni corrected; Fig. 5A,C). For example, the first component could be read out with significantly above-chance accuracy in both languages; see below for a more detailed discussion of which factors could be decoded. Note that confidence intervals were wider for German, which we attribute to the much smaller number of trials in the German experiment. Time-resolved decoding (Figure 5B,D) indicated that information about these factors was recoverable from brain activity beginning at ∼250 msec and peaking around 350-400 msec, i.e., again in the time window of the N400, the best-established semantic ERP component.

**Figure 5:**
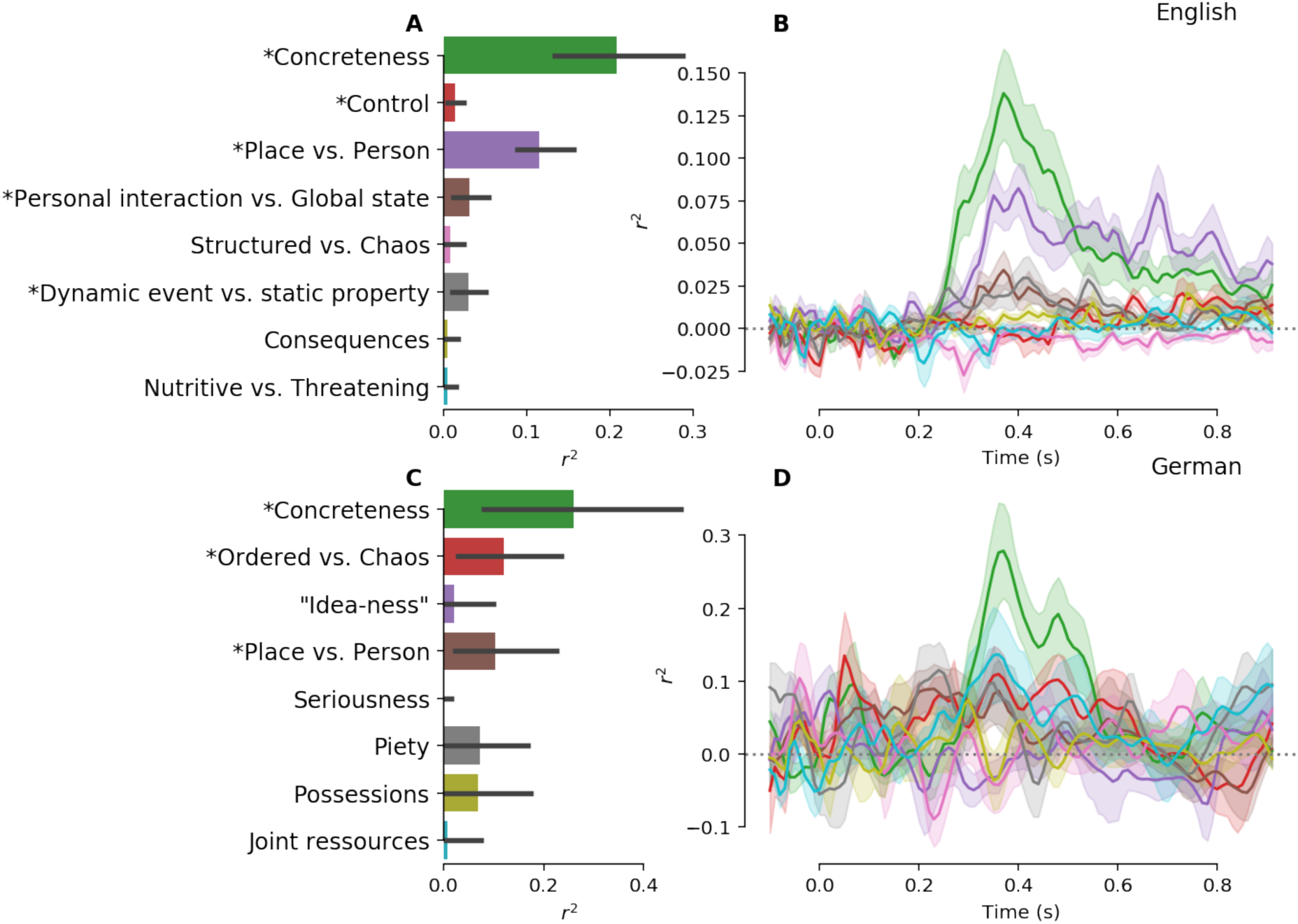
Decoding semantic dimensions from EEG data. A: Prediction accuracy (measured as the squared, signed correlations between predicted and actual values) for the prediction of eight vector-space factors from EEG activity between 300 - 500 msec, for English data. Labels on the ; axis reflect a subjective interpretation of the factor meanings (see Methods and Table 1). Error bars represent the 99.375% (95% corrected for multiple comparisons) bootstrapped confidence intervals; asterisks indicate decoding accuracies significantly different from chance. B: Time-resolved decoding scores for the same eight factors, when repeating the prediction at each sample time point of the EEG signal. Colors as on left plot. Shaded outlines reflect 68% bootstrapped confidence intervals over folds (uncorrected). C/D: Same, but for the German dataset (note the different scaling of the y axis in D as compared to B).

**Table 1:**
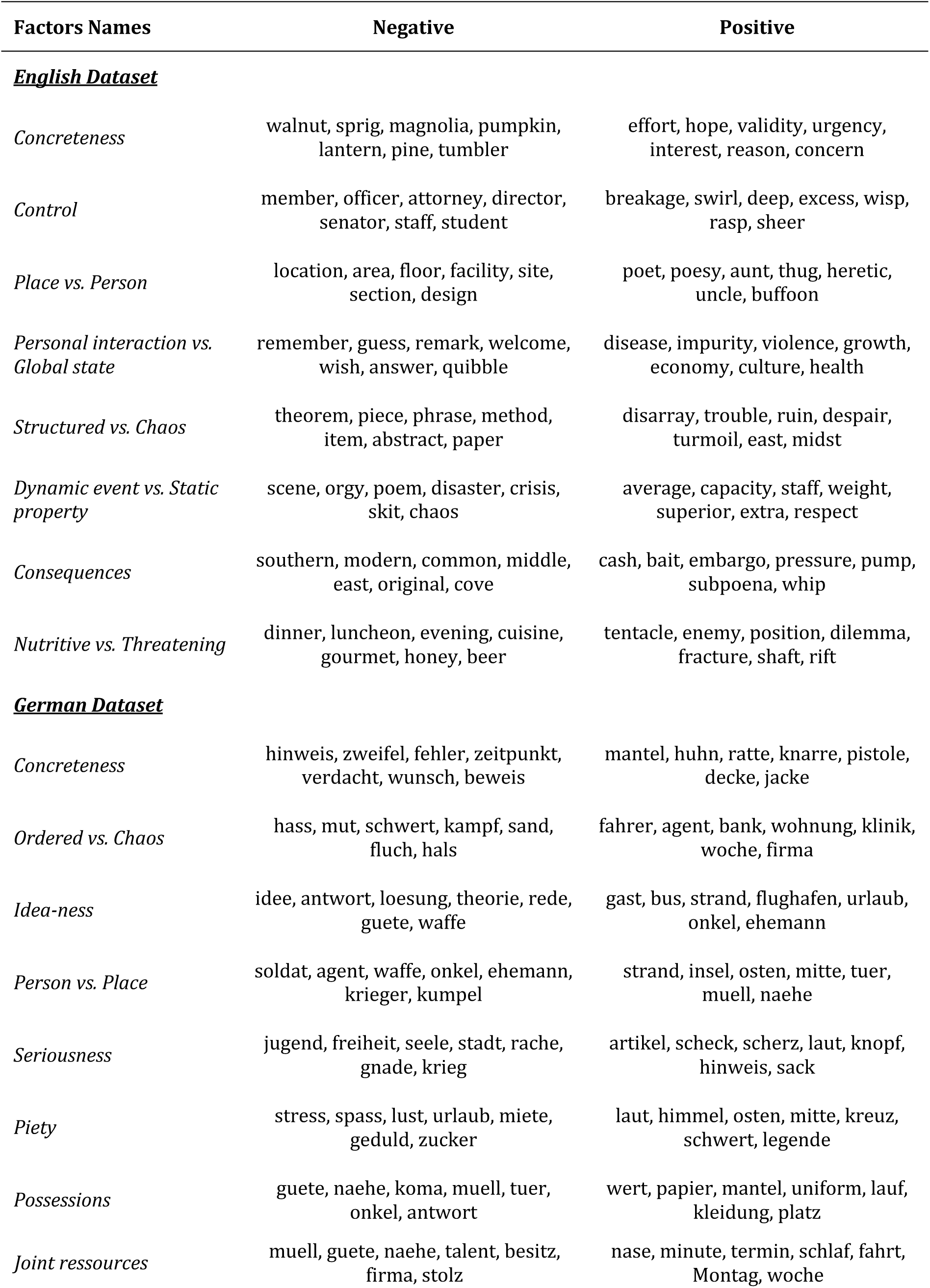
Tentative labels for each component derived from the dimensionality reduction of word vector space, plus most high/low-scoring words. Note that words are shown as they were fed to the word vector model (i.e., all lower cased). The seven most strongly negatively and most strongly positively scoring words are shown as example.

### 3.3 Kernel PCA Factor Structure

The first eight factors identified via Kernel PCA decomposition in the distributional models for each language are shown in Table 1. The semantic interpretation of these factors (provided by trained linguists; see Methods for details) is ad-hoc, is not constrained by brain activity, and does not conclusively establish that these factors are intrinsic to the structure of the distributional models. Nevertheless, it is noteworthy that the first of these factors in both languages constituted one of the most fundamental dimensions of semantics discussed in the literature, i.e., Concreteness, and that at least two other factors (Place vs. Person; Order/Structure vs. Chaos) appear in both languages. Even more importantly, these clearly semantic components could be read out from brain activity: We found significantly above-chance decoding scores for Concreteness (German and English Factor 1), Place vs. Person (English Factor 3, German Factor 4), and Ordered vs. Chaos (German Factor 2). In addition to these factors, decoding scores significantly (p < .05, Bonferroni corrected) exceeded chance for Factors 4 and 6 (English); see Figure 5A, C.

Again, we wish to emphasize here that the observed factor structure is in no way indicative of a true semantic feature space that we assume to be represented in the human brain, and choosing different analysis parameters could lead to very different factor structures. This analysis, thus, is merely a tool for verifying certain aspects of our methodology, intended to demonstrate that at least some of the predictive power of distributional models on ERP data stems from the genuinely semantic information they represent.

### 3.4 Visualising decoding and encoding patterns

By visualizing the spatial structure of these effects (see Fig. 6), we find, very broadly speaking, that the spatial distribution of encoding performance as well as the informativity of electrodes for the decoding procedure partially, but not fully, overlap with the scalp distribution of the N400 effect known from other studies of semantic processing (Kutas & Federmeier, 2011). I.e., we observe that encoding performance is highest at central electrodes near the midline, and that for at least 4 Kernel PCA components (1, 5, 7, and 8), the most important predictors in the decoding analysis are similarly found at central electrodes near the midline. However, other components are better predicted by other electrodes, perhaps indicating that different neural generators are involved in these representations.

**Figure 6:**
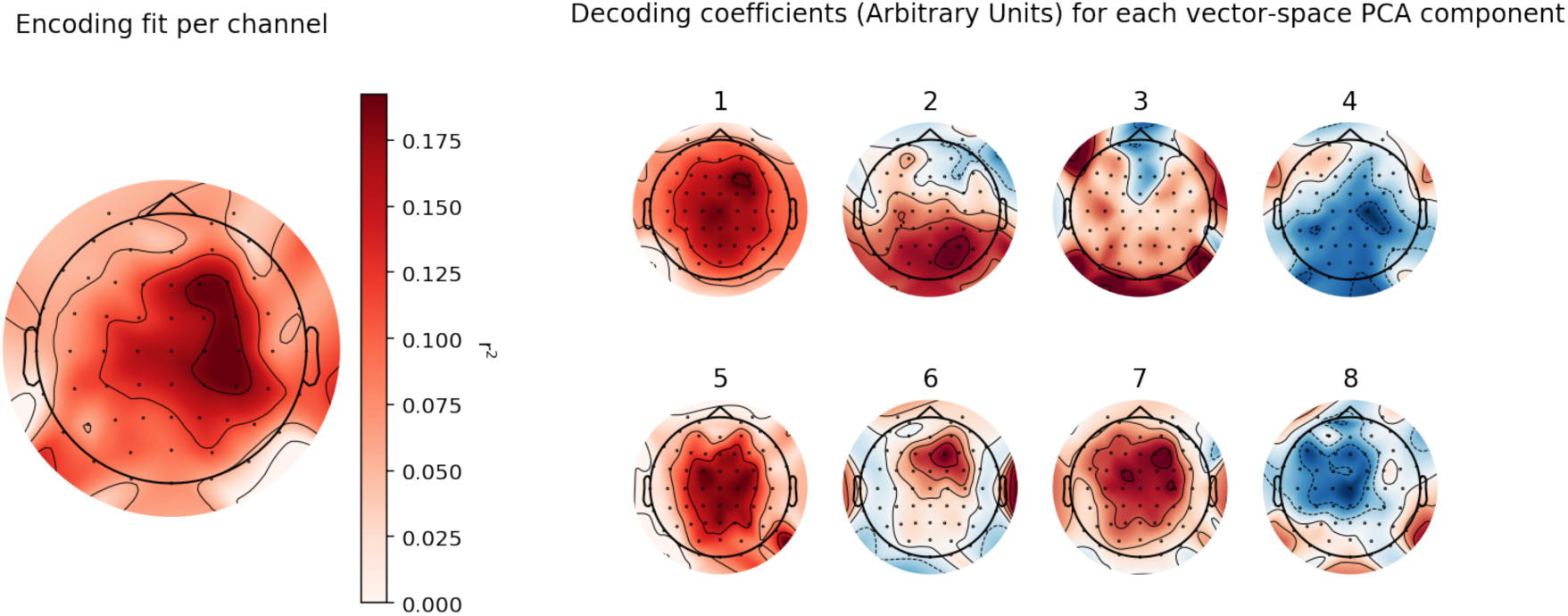
Visualization of the scalp distribution of encoding and decoding models. Left: Scalp distribution map showing the r^2 of the encoding model, for the N400 time window (300 – 500 msec), per electrode. Right: For each of the 8 kernel PCA components, the variance-normalized coefficients of the respective decoder (i.e., the forward model) are shown per electrode. I.e., more red/blue coloring indicates that a more positive/negative going ERP leads to a higher prediction on the respective kernel PCA score (German dataset; compare Figure 5). These values are dimensionless.

## 4. Discussion

In this study, we have demonstrated that brain activity measured with EEG (more specifically, the distribution of ERP timecourses over the scalp) encodes distributional vector representations of word meaning, as derived from nowadays well established prediction-based distributional models. This result suggests that semantic vector spaces represent semantic relationships between words in a manner that is at least partly shared with how the brain represents word semantics. More specifically, words that occupy adjacent spaces in the high-dimensional vector space representations of these models also induce similar brain activation states. These correspondences were largely the same for two experiments conducted in different languages (i.e., English and German) and requiring two different behavioral tasks. Also, our results did not change substantially when using a number of different implementations of distributional models.

On the one hand, our results indicate that distributional vector embeddings of word meaning allow accurate simulations of brain states (i.e., the *encoding* of word vectors in brain activity), indicating that word semantics, as expressed in word vectors, entrain brain states into predictable patterns. On the other hand, our data also demonstrate that vector-based word embeddings allow reading out (i.e., *decoding*) parts of the meaning of words from brain activity. This indicates that a semantic architecture at least partially similar to distributional word vector spaces is encoded in brain activity – even though these may not be directly interpretable semantic features. It is particularly noteworthy that the vector-space components which could be read out (decoded) from brain activity were independently described by trained linguists as corresponding to axes of meaning (rather than nonsemantic aspects of word knowledge, such as a word’s length or frequency of occurrence). This is indeed expected if brain activity elicited during word processing contains representational information about word meaning and if the nature of this neural representation is at least partly related to the distributional nature of word vectors.

In the following, we will discuss the implications of these results for the cognitive neuroscience of meaning, i.e., for understanding the how, what, and when of representing the semantics of words in the human mind and brain. First, we will discuss how the present results relate to various models of semantic representation. Then, we will argue that some distinctions between these models (e.g., between feature- and exemplar-based architectures) are irrelevant, so that the core difference between models is which feature space they (perhaps implicitly) operate in. We will also review in somewhat more detail Jerry Fodor’s argument that *conceptual* feature spaces – unlike the not directly interpretable distributional word vector models we studied here – are implausible because they are circular (Fodor & LePore, 1999). From these considerations, combined with our empirical results, we will elaborate that distributional models – like those whose fit to brain activity we tested here – are reasonable contenders for the architecture of the semantic system. Lastly, we connect our results to previous findings and theories from the cognitive neurosciences, and argue that brain activity in the N400 time window reflects not only retrieval/integration-related *processing*, but also (context-independent) semantic *representations*.

### 4.1 Models of semantics representation

What options for models of semantic representations are there? Distributional models like word2vec have been proposed only recently, joining a long list of alternative proposals. *Empiricist state-space* theories (e.g., Churchland and Sejnowski, 1990) propose that word meaning can be understood as the ranking of a word on various experiential dimensions, i.e., categories derived from our senses. An important role of not only the sensory, but also the motor system for the representation of semantics is suggested by *Embodiment theories* (e.g., Gallese and Lakoff, 2005; Pulvermüller, 2013). These theories claim that we represent both concrete and abstract categories with reference to our sensory and motor systems. There are also *nativist feature theories*, which propose that our mental concepts can be decomposed, although these features are not understood as experiential in nature, but innate (e.g., Pinker, 2008). Finally, *Exemplar or Prototype theories* (e.g., Edelman, 1995; Rosch, 1999) postulate that concepts are represented as, or with respect to, exemplars. Although prototypes must not necessarily correspond to objects or persons we have seen in the real world with our own eyes, they are constructed based on such experiential phenomena. Assigning a specific token to a semantic concept then corresponds to checking how similar it is to the respective prototype.

Under all these approaches, semantic representations can be conceptualized in matrix form, i.e., geometrically as vectors or as points in high-dimensional state spaces (e.g., Churchland, 1993; Kriegeskorte & Kievit, 2013; Warglien & Gärdenfors, 2013). The disagreement between these models is about what the axes of the space or matrix correspond to: Nativist models assume they reflect innate, abstract protosemantic concepts (as those in Dowty, 1994; see also Pinker, 2008). Empiricist (Churchland, 1993; Prinz, 2004) as well as embodiment models (Pulvermüller, 2013) suggest experiential features, whereas in prototype/exemplar models (Edelman, 1995) exemplars take the role of the features – i.e., a concept’s position is described with regards to its distance to other concepts.

Interestingly, it has turned out to be irrelevant if these features are categorical or continuous (+ANIMATE vs. DEGREE OF ANIMACY; cf. Tissier et al., 2018) in nature. Similarly, it turns out to be irrelevant if distances are represented with regards to other concepts (i.e., exemplars or prototypes) or to features. Conceptually, this can be seen by considering that a distance between two items corresponds to a consideration of feature overlap – in other words, the distance between two exemplars or prototypes implicitly refers to a feature space. If a different feature space is chosen, new distances result. Thus, exemplar models operate at least implicitly in a feature space, which has mathematically been formalized in the form of the so-called Representer Theorem (Schölkopf, Herbich & Smola, 2001; Schölkopf et al., 2007; for a related development, see Nunez-Elizalde et al., 2019). This finding establishes that for a broad class of inference processes, the so-called primal and dual problems (in this case corresponding to feature- vs. exemplar-based encoding processes) are equivalent; conducting the inference in one space gives the same results as in the other (see also Fig. 1, bottom). Thus, any query of the semantic model yields the same results for a feature-based representation as compared to a representation in an exemplar space defined by the same features. Moving from feature- to exemplar space simply requires applying a *kernel function* (i.e., a similarity measure, yielding the distance between entities in feature space; this inspired our choice of Kernel PCA in the decoding section). This implies for a quantitative evaluation of alternative models of semantic representations, that the crucial question does not pertain to the general architecture of the model – i.e., a featural vs. exemplar-based architecture. Rather, it is critical to understand of what kind the features are that define the semantic state space. We hoped to elucidate on this question by testing how well various feature spaces can predict brain activity elicited during word perception.

We found that distributional word vector models allow the prediction (*encoding*) of human brain activity, and that this prediction is significantly better than prediction based on a classical taxonomic baseline model. This superiority of distributional vector-based representations to taxonomic models suggests that logical relations such as “x is a y” (hyponymy/hyperonymy) are not sufficient to grasp the neural implementation of semantic coding. Richer associations – perhaps including dimensions that resist verbal descriptions – are superior in explaining brain activity, and are in turn encoded in brain activity. In this sense, our work complements previous research on feature-based explanations of meaning representation (Fyshe, Sudre, Wehbe, Murphy, & Mitchell, 2012; Simmons, Hamann, Harenski, Hu, & Barsalou, 2008). In this sense, our work converges with the encoding of BOLD responses to continuous, naturalistic language stimulation reportedby Huth et al. (2016) but extends this work by demonstrating a systematic relationship between brain activation during isolated (i.e., context-free) word reading and distributional word vectors.

On the other hand, our *decoding* results suggest that the neural activity explained by distributional word vector models is at least in part genuinely semantic in nature. Some of the features we can decode are unsurprising; for example, concreteness is known to be a major determinant of lexical representation and was shown to modulate brain activity (Dufau et al., 2015). Others can be interpreted as evolutionarily relevant signals – i.e., threat, food, or social aspects of meaning, which have been independently noted to affect word processing (Bentall & Kaney, 1989). However, the mere observation that a feature can be decoded from brain activity does not in itself mean that it is a major axis of the neural representation of meaning. The encoding process is bottlenecked on both sides, by the quality of the vectorial representations (i.e., it is possible that a highly important dimension simply cannot be learned on data, or by models as those we use here), and by the signal to noise ratio of the brain data. Our dimensionality reduction of the word vectors was entirely based on patterns of (co)variance within the word vectors themselves, not on what information is encoded in the neural signal. Put differently, the decoding procedure can only discover statistical relations between brain activity and semantic patterns identified independently of brain activity. Thus, while our results are suggestive of distributional semantics, they do not provide any finer insights into the specifics of human semantic representations.

Interestingly, there are also indications that distributional, word2vec-class models can also yield inferior results compared to alternative models. For example, Huth et al. 2016 performed a supplementary analysis comparing their primary analysis to a Word2Vec-based distributional model. Contrary to our results, these authors found that Word2Vec explained fMRI activity above chance, but significantly worse than their primary model. The most obvious differences between these two studies are the recording modality (fMRI vs. EEG) and the stimulus coherence (narratives in Huth et al., 2016, as opposed to single words in our study). One important difference between EEG/ERP and fMRI is that fMRI recordings have superior signal-to-noise levels. This suggests that differences in SNR per se cannot explain why Word2Vec performed better in our EEG/ERP study. A major difference is furthermore the superior temporal resolution of EEG. We thus speculate that that the high temporal resolution of EEG/ERP may have allowed us to specifically identify the moment in time where processing of word meaning is performed. In contrast, fMRI conflates different cognitive processes into one datapoint, which may reduce its ability to capture the temporally circumscribed instantiation of meaning during word comprehension.

Another difference between the materials in our study and this previous work concerns the ongoing integration of words into phrasal and sentence contexts – contexts which were available to participants with the materials employed by Huth et al., 2016, but not with our single-word materials. As many or even all words are polysemuous (Hagoort et al., 2004), many of our stimulus words were ambiguous between multiple meanings, and some even between word classes (e.g., *fish*, which can function both as noun or verb). In contrast, word processing with constraining contexts is a different task where only a narrow subset of the possible semantics associated with a word are evoked in order to possibly be integrated into the mental representation of the sentence. Moreover, continuous narratives consist of both content and function words, many of which carry their own semantics. This might lead to important differences in processing between continuous narratives vs. individual words. However, while this crucial difference may underlie differences between our findings and the results reported by Huth et al. (2016), the direction of the difference is surprising because intuitively, one might have expected to find the context-dependent Word2Vec analysis to perform superior for words presented in context. In the long run, theoretical work will have to develop a theory that can simultaneously accommodate both the integration of meaning in context, as well as the representation of meaning evoked by individual words presented in isolation.

We also note that we found vectors based on the original Word2Vec implementation (also used by Huth et al., 2016) substantially inferior to the improved FastText-based vectors we employed. It is thus possible that improved distributional or taxonomic models will both yield new results. Lastly, we would also like to point out that differences in task and material are highly likely to determine which feature spaces prove superior in different study contexts. For example, neural signatures of taxonomic judgement tasks are likely to favor taxonomic models. Future work will thus also have to explicitly explore the effect of such ‘design’ variables on the encoding and decoding results.

Lastly, it should be noted that the specific stimuli employed may have a strong impact on our results (as well as on other studies using similar model-based approaches): The extent to which results can be generalized beyond the given study is limited by the specific nature of the stimuli – which here consisted of short nouns and, for the German data, of clearly abstract or concrete words. For example, as an inherent limitation of distributional models, they capture much less well the meaning of very low-frequency words. Due to the length restrictions of our experiment, we did not employ extremely long and rare words such as “heteroskedasticity”, for which a distributional model - based fit is expected to be worse. Similarly, little can be said about, e.g., regular compounds.

### 4.2 Circularity of conceptual features, and the appeal of distributional models

In the literature on semantic architectures and representations, we have so far not found an example of a comprehensive list of semantic features – in the sense of a feature list that could clearly capture the semantics of any non-trivial set of word meanings. To the best of our knowledge, there is no such canonical list of semantic features with a claim to completeness. Put differently, theories of semantics (see above) are proposals regarding the *kind* of features that define the representation of semantic meaning, but none of these models has so far spelled out testable proposals for these features. We argue that this hinders the empirical evaluation of the proposed theoretical models, as these models cannot generate quantitative and thus testable predictions without a clear definition of the involved features (i.e., without an explicit enumeration of the labels of the axes of the feature space).

Fodor and LePore (1999) and Fodor (1970) have argued that any enumeration of specific features would be futile, because any definition of concepts with features that are themselves conceptual in nature is either insufficient or trivial. For example, it would be unsatisfactory to define *cat* as having a feature +CAT. By this circular definition, the set of features would equal or exceed the number of word meanings. Instead, feature semantics strives to achieve a parsimonious description by identifying redundancies. Such an approach seems to work well for some showcase concepts such as *bachelor* (+MALE, +UNMARRIED). On a closer look, however, most words resist a decomposition into any convincing bundle of features as long as they do not become tautological. For example, *die* means BECOME DEAD, *dead* means HAVE DIED, etc. (Fodor, 1970). Similarly, *cat* cannot be defined by any parsimonious set of features excluding the feature +CAT. Thus, a model of semantic representation where the discriminating features (also called axes or dimensions) are concept-like is – on virtue of circularity – implausible.

Already in 1997, Landauer and Dumais proposed that a practical approach (called Latent Semantic Analysis, or LSA) for deriving vectorial representations of words is to enumerate what contexts (e.g., documents, sentences …) words occur in, and then identify latent variables underlying this massive *contexts* × *words* matrix. While LSA is surprisingly effective, it is unlikely that this is similar to how humans actually learn; toddlers probably do not have the capacity to verbatim store thousands of paragraphs. Models like word2vec make distributional vectors *learnable* by implicitly connecting them to predictive processing – i.e., a process human brains are well known to be constantly engaged in (e.g., Clark, 2013). This offers a promising improvement on earlier models. The important aspect of word2vec and LSA-style models, relative to classical conceptual-feature space models, is that their axis labels are not conceptual, which allows them to evade the circularity critique. By this, we suggest, they become surprisingly plausible candidates for the representational architectures of the human mind. They also have the benefit of being already available in implemented form – unlike competing models, which are often only specified conceptually. This, in turn, opens up a framework for robust and quantitative tests of theories of semantic representation. The data we present in the present study provide a natural extension of such conceptual considerations by empirically demonstrating the relationship between word vectors and brain states.

Proponents of grounded or embodied semantics (for a recent review, see Hauk & Tschentscher, 2013) would argue that the dimensions of representation of distributed models lack a biological grounding. However, these two approaches may in fact be surprisingly complementary. For example, Frome et al. (2013) have presented work indicating that combining perceptual and (text-based) distributional information may be a key aspect to rapid word acquisition (in their case demonstrating inference of semantics based on both word distributions and images). This suggests that combining statistical information about word context with the rich understanding of the physical world typical of human learners may be an essential aspect for improving the performance of distributional models to reach human-equivalent levels. For this reason, we prefer to see current empiricist distributional models (like investigated in the present work) not as an alternative to grounded theories of semantics. Instead, we think that they supplement each other, and integrating such perspectives could be a key step in furthering our understanding of human semantic knowledge.

### 4.3 Meaning and the N400 time window

Our finding that semantic vector space information is encoded in brain activity measured between 300 and 500 ms after word onset has interesting implications for understanding the cognitive and neural processes underlying activity in the time window of the N400 component of the event-related brain potential. The N400 has in a large body of empirical research been shown to covary with the difficulty of semantic processing of words. In particular, it has been linked to lexical access and the integration of a word into a sentence context (Kutas & Federmeier, 2000; Kutas & Federmeier, 2011). Here, we find that neurophysiological brain activity in the N400-time window contains information that is partially structured akin to distributional semantic models – in a situation without a context to integrate words into. This indicates that N400-time window activity reflects more than the difficulty of cognitive processes associated with retrieving word meaning and/or integrating it into the current context – which is the currently most widely accepted interpretation of this ERP effect (Lau et al., 2008; Kutas & Federmeier, 2011). Such a gradient of context-dependent processing difficulty is also mainly what previous attempts to encode word vectors in EEG activity have observed (Broderick et al., 2017; Ettinger et al., 2016). Instead, our findings indicate that EEG activity in the N400 time window also directly reflects a word’s meaning. This is not surprising; after all, one could even go so far as to say that the very purpose of words is to systematically shift the brain of the perceiver into certain states that represent their meaning. Thus, there *should* be a correspondence between the meaning of words perceived in isolation and the perceiver’s brain states. However, this was so far not directly accessible to typical empirical neurocognitive studies, as most previous research has focused on context-dependent *processes* rather than context-independent *representations*.

The observed encoding of semantic vector spaces in brain activity opens up a novel set of questions for researchers investigating the brain bases of meaning and language. Ideally, cognitive semantics and research on the neurobiology of language can jointly address the question of how words, after they have been processed (accessed, retrieved, integrated, …), are represented in brain activity. On the other hand, our results also suggest novel ways of investigating the corresponding cognitive architecture. For example, the encoding approach chosen here allows transcending factorial condition-contrast experiments, instead making it possible to directly compare representational frameworks with each other (Kriegeskorte & Kievit, 2013). Integrating our data with the currently available body of evidence indicates that processing-dependent aspects of semantics play out in approximately the same time window as representation-dependent aspects (i.e., roughly the N400 time window), suggesting that there may not be a strict boundary between representing and processing – as suggested by, e.g., dynamicist models of semantic processing (Elman, 2009; Kutas & Federmeier, 2011; Lupyan & Lewis, 2017).

### 4.4 Outlook

Distributional word vectors (learned from statistical patterns in texts) provide a demonstration of how a representational space sufficient to afford human semantic cognition could work. An important implication of our finding that semantic representations are encoded in brain activity (i.e., the event-related brain potential), and thus can also be decoded from EEG activity, is that *any* theory of semantics can be tested against neural data, to the extent that it can be quantified. I.e., any independently motivated list of features could easily be encoded and the result compared to a suitable benchmark (such as the models tested here). Our results constitute one such benchmark; a research strategy for competing models should be to aim at producing higher portions of explained variance than reported here, i.e., to beat the prediction accuracy obtained from distributional word vectors in the present work.

But is it plausible to assume that the brain computes a Word2Vec-like analysis when learning words, or that it represents its semantic knowledge in abstract multidimensional vector form? Most certainly not. Existing methods, while powerful, are far from human-like in performance, even if they are trained on much larger corpora than children need for learning. And most models – including those tested here – are unimodal; i.e., they do not derive any support from sensory experiences, generalization from other domains, or any cognitive priors, which would be a highly inefficient strategy for brains to use. We do not wish to suggest that mental semantics literally equal the Word2Vec algorithm or similar distributional models. However, we agree with Landauer and Dumais (1997; see above) that distributional word vector models avoid certain key problems of alternative approaches, most importantly the circularity critique developed by Fodor (1996). Combined with the successes of distributional models in de- and encoding of brain activity, they establish a strong baseline for the modelling of word semantics-associated brain activity, which competing models must beat.

## Acknowledgements

The authors wish to thank Jonathan Grainger and Stephane Dufau for making available the English EEG dataset. Edvard Heikel organized collection of the German EEG data. The MNE-Python Encoding team, in particular Chris Holdgraf, Eric Larson, Alexandre Gramfort, Denis Engemann, and Jean-Remi King, helped with analysis code and conceptual discussion.

## References

Aziz-Zadeh, L., Fiebach, C. J., Naranayan, S., Feldman, J., Dodge, E., & Ivry, R. B. (2008). Modulation of the FFA and PPA by language related to faces and places. Social Neuroscience, 3(3-4), 229–238.

Aziz-Zadeh, L., & Damasio, A. (2008). Embodied semantics for actions: Findings from functional brain imaging. Journal of Physiology-Paris, 102(1-3), 35–39.

Baroni, M., Dinu, G., & Kruszewski, G. (2014). Don’t count, predict! A systematic comparison of context-counting vs. context-predicting semantic vectors. In Proceedings of the 52nd Annual Meeting of the Association for Computational Linguistics (Volume 1: Long Papers) (Vol. 1, pp. 238–247).

Bentall, R. P., & Kaney, S. (1989). Content specific information processing and persecutory delusions: An investigation using the emotional stroop test. British Journal of Medical Psychology, 62(4), 355–364.

Bojanowski, P., Grave, E., Joulin, A., & Mikolov, T. (2016). Enriching word vectors with subword information. CoRR, abs/1607.04606. Retrieved from http://arxiv.org/abs/1607.04606

Borghesani, V., & Piazza, M. (2017). The neuro-cognitive representations of symbols: The case of concrete words. Neuropsychologia, 105, 4–17.

Broderick, M. P., Anderson, A. J., Di Liberto, G. M., Crosse, M. J., & Lalor, E. C. (2018). Electrophysiological Correlates of Semantic Dissimilarity Reflect the Comprehension of Natural, Narrative Speech. Current Biology, 28(5), 803–809.e3.

Brysbaert, M., Buchmeier, M., Conrad, M., Jacobs, A. M., Bölte, J., & Böhl, A. (2011). The word frequency effect. Experimental Psychology, 58(5), 412–424.

Churchland, P. M. (1993). State-space semantics and meaning holism. Philosophy and Phenomenological Research, 53(3), 667–672. Retrieved from http://www.jstor.org/stable/2108090

Churchland, P. S., & Sejnowski, T. J. (1990). Neural representation and neural computation. Philosophical Perspectives, 4, 343–382.

Clark, A. (2013). Whatever next? Predictive brains, situated agents, and the future of cognitive science. Behavioral and brain sciences, 36(3), 181–204.

Dowty, D. (1994). The role of negative polarity and concord marking in natural language reasoning. In Semantics and Linguistic Theory (Vol. 4, pp. 114–144).

Dufau, S., Grainger, J., Midgley, K. J., & Holcomb, P. J. (2015). A thousand words are worth a picture: Snapshots of printed-word processing in an event-related potential megastudy. Psychological Science, 26(12), 1887–1897.

Edelman, S. (1995). Representation, similarity, and the chorus of prototypes. Minds and Machines, 5(1), 45–68. https://doi.org/10.1007/BF00974189

Elman, J. L. (1991). Distributed representations, simple recurrent networks, and grammatical structure. Machine Learning, 7(2-3), 195–225.

Elman, J. L. (2009). On the meaning of words and dinosaur bones: Lexical knowledge without a lexicon. Cognitive Science, 33(4), 547–582.

Ettinger, A., Feldman, N., Resnik, P., & Phillips, C. (2016). Modeling n400 amplitude using vector space models of word representation. In A. Papafragou, D. Grodner, D. Mirman, J.C. Trueswell (Eds.), Proceedings of the 38th Annual Conference of the Cognitive Science Society (pp. 1445–1450). Austin, TX: Cognitive Science Society.

Felbo, B., Mislove, A., Søgaard, A., Rahwan, I., & Lehmann, S. (2017). Using millions of emoji occurrences to learn any-domain representations for detecting sentiment, emotion and sarcasm. arXiv preprint arXiv:1708.00524.

Fiebach, C. J., & Friederici, A. D. (2004). Processing concrete words: fMRI evidence against a specific right-hemisphere involvement. Neuropsychologia, 42(1), 62–70.

Fodor, J. A. (1970). Three Reasons for Not Deriving “Kill” from “Cause to Die”. Linguistic Inquiry, 1(4), 429–438.

Fodor, J. A., & LePore, E. (1999). All at sea in semantic space: Churchland on meaning similarity. The Journal of Philosophy, 96(8), 381–403.

Fodor, J. (2004). Having concepts: A brief refutation of the twentieth century. Mind & Language, 19(1), 29–47.

Fyshe, A., Sudre, G., Wehbe, L., Murphy, B., & Mitchell, T. (2012). Decoding Word Semantics from Magnetoencephalography Time Series Transformations. Proceedings of the 3rd Workshop on Machine Learning and Inference in Neuroimaging, NIPS.

Gauthier, J., & Ivanova, A. (2018). Does the brain represent words? An evaluation of brain decoding studies of language understanding. arXiv preprint arXiv:1806.00591.

Gallese, V., & Lakoff, G. (2005). The brain’s concepts: The role of the sensory-motor system in conceptual knowledge. Cognitive Neuropsychology, 22(3-4), 455–479.

Gramfort, A., Luessi, M., Larson, E., Engemann, D. A., Strohmeier, D., Brodbeck, C., Goj, R., Jas, M., Brooks, T., Parkkonen, L., … Hämäläinen, M. (2013). MEG and EEG data analysis with MNE-Python. Frontiers in neuroscience, 7, 267. doi:10.3389/fnins.2013.00267

Günther, F., Dudschig, C., & Kaup, B. (2016a). Latent semantic analysis cosines as a cognitive similarity measure: Evidence from priming studies. The Quarterly Journal of Experimental Psychology, 69(4), 626–653.

Günther, F., Dudschig, C., & Kaup, B. (2016b). Predicting lexical priming effects from distributional semantic similarities: A replication with extension. Frontiers in Psychology, 7, 1646.

Hagoort, P., Hald, L., Bastiaansen, M. C. M., & Petersson, K.-M. (2004). Integration of word meaning and world knowledge in language comprehension. Science, 304(5669), 438– 441.

Hamp, B., & Feldweg, H. (1997). GermaNet - a Lexical-Semantic Net for German. Proceedings of ACL Workshop Automatic Information Extraction and Building of Lexical Semantic Resources for NLP Applications.

Hastie, T., Tibshirani, R., & Friedman, J. H. (2009). The Elements of Statistical Learning. New York: Springer.

Hauk, O., Johnsrude, I., & Pulvermüller, F. (2004). Somatotopic representation of action words in human motor and premotor cortex. Neuron, 41(2), 301–307.

Hauk, O., & Tschentscher, N. (2013). The Body of Evidence: What Can Neuroscience Tell Us about Embodied Semantics?. Frontiers in psychology, 4, 50. doi:10.3389/fpsyg.2013.00050

Holdgraf, C. R., Rieger, J. W., Micheli, C., Martin, S., Knight, R. T., & Theunissen, F. E. (2017). Encoding and decoding models in cognitive electrophysiology. Frontiers in Systems Neuroscience, 11, 61. https://doi.org/10.3389/fnsys.2017.00061

Huth, A. G., de Heer, W. A., Griffiths, T. L., Theunissen, F. E., & Gallant, J. L. (2016). Natural speech reveals the semantic maps that tile human cerebral cortex. Nature, 532(7600), 453–458.

Jäkel, F., Schölkopf, B., & Wichmann, F. A. (2007). A tutorial on kernel methods for categorization. Journal of Mathematical Psychology, 51(6), 343–358.

Jung, T.-P., Makeig, S., Humphries, C., Lee, T. W., McKeown, M. J., Iragui, V., & Sejnowski, T. J. (2000). Removing electroencephalographic artifacts by blind source separation. Psychophysiology, 37(2), 163–178.

Jurafsky, D., & Martin, J. H. (2014). Speech and language processing (Vol. 3). London: Pearson.

Ju, R., Zhou, P., Li, C. H., & Liu, L. (2015, October). An efficient method for document categorization based on word2vec and latent semantic analysis. In 2015 IEEE International Conference on Computer and Information Technology; Ubiquitous Computing and Communications; Dependable, Autonomic and Secure Computing; Pervasive Intelligence and Computing (pp. 2276–2283). IEEE.

Kemmerer, D., Castillo, J. G., Talavage, T., Patterson, S., & Wiley, C. (2008). Neuroanatomical distribution of five semantic components of verbs: Evidence from fMRI. Brain and Language, 107(1), 16–43.

King, J. R., & Dehaene, S. (2014). Characterizing the dynamics of mental representations: the temporal generalization method. Trends in cognitive sciences, 18(4), 203–210.

King, J. R., Gwilliams, L., Holdgraf, C., Sassenhagen, J., Barachant, A., Engemann, D., … & Gramfort, A. (2018). Encoding and Decoding Neuronal Dynamics: Methodological Framework to Uncover the Algorithms of Cognition.

Krause, C. M., Aström, T., Karrasch, M., Laine, M., & Sillanmäki, L. (1999). Cortical activation related to auditory semantic matching of concrete versus abstract words. Clinical Neurophysiology: Official Journal of the International Federation of Clinical Neurophysiology, 110(8), 1371–1377.

Kriegeskorte, N., & Kievit, R. A. (2013). Representational geometry: integrating cognition, computation, and the brain. Trends in cognitive sciences, 17(8), 401–412.

Kutas, M., & Federmeier, K. D. (2000). Electrophysiology reveals semantic memory use in language comprehension. Trends in Cognitive Science, 4(12), 463–470.

Kutas, M., & Federmeier, K. D. (2011). Thirty years and counting: finding meaning in the N400 component of the event-related brain potential (ERP). Annual Review of Psychology, 62, 621–647.

Lambon Ralph, M. A., & Patterson, K. (2008). Generalization and differentiation in semantic memory: insights from semantic dementia. Annals of the New York Academy of Sciences, 1124(1), 61–76.

Landauer, T. K., & Dumais, S. T. (1997). A solution to Plato’s problem: The latent semantic analysis theory of acquisition, induction, and representation of knowledge. Psychological Review, 104(2), 211–240.

Lau, E. F., Phillips, C., & Poeppel, D. (2008). A cortical network for semantics:(de) constructing the N400. Nature Reviews Neuroscience, 9(12), 920.

Levy, O., & Goldberg, Y. (2014). Neural Word Embedding as Implicit Matrix Factorization. In Z. Ghahramani, M. Welling, C. Cortes, N. D. Lawrence, & K. Q. Weinberger (Eds.), Advances in Neural Information Processing Systems 27 (pp. 2177– 2185). Curran Associates, Inc. Retrieved from http://papers.nips.cc/paper/5477-neural-word-embedding-as-implicit-matrix-factorization.pdf

Lupyan, G., & Lewis, M. (2017). From words-as-mappings to words-as-cues: The role of language in semantic knowledge. Language, Cognition and Neuroscience. Retrieved from https://doi.org/10.1080/23273798.2017.1404114

Mandera, P., Keuleers, E., & Brysbaert, M. (2017). Explaining human performance in psycholinguistic tasks with models of semantic similarity based on prediction and counting: A review and empirical validation. Journal of Memory and Language, 92, 57– 78.

Mikolov, T., Chen, K., Corrado, G.S., & Dean, J. (2013). Efficient Estimation of Word Representations in Vector Space. Proceedings of Workshop at ICLR, 2013.

Mikolov, Tomas, Le, Quoc V., and Sutskever, Ilya (2013b). Exploiting Similarities among Languages for Machine Translation. arXiv:1309.4168

Mikolov, T., Grave, E., Bojanowski, P., Puhrsch, C., & Joulin, A. (2017). Advances in pre-training distributed word representations. arXiv Preprint arXiv:1712.09405.

Miller, G. A. (1995). WordNet: A lexical database for English. Communications of the ACM, 38(11), 39–41.

Mitchell, T. M., Shinkareva, S. V., Carlson, A., Chang, K. M., Malave, V. L., Mason, R. A., & Just, M. A. (2008). Predicting Human Brain Activity Associated with the Meanings of Nouns. Science, 320(5880), 1191–1195.

Nunez-Elizalde, A. O., Huth, A. G., & Gallant, J. L. (2019). Voxelwise encoding models with non-spherical multivariate normal priors. NeuroImage, 197, 482–492.

Patterson, K., Nestor, P. J., & Rogers, T. T. (2007). Where do you know what you know? The representation of semantic knowledge in the human brain. Nature Reviews Neuroscience, 8(12), 976.

Pedregosa, F., Varoquaux, G., Gramfort, A., Michel, V., Thirion, B., Grisel, O., … Dubourg, V. (2011). Scikit-learn: Machine learning in Python. The Journal of Machine Learning Research, 12, 2825–2830.

Pennington, J., Socher, R., & Manning, C. (2014). Glove: Global vectors for word representation. In Proceedings of the 2014 conference on empirical methods in natural language processing (EMNLP) (pp. 1532–1543).

Pereira, F., Lou, B., Pritchett, B., Ritter, S., Gershman, S. J., Kanwisher, N., … & Fedorenko, E. (2018). Toward a universal decoder of linguistic meaning from brain activation. Nature communications, 9(1), 963.

Pinker, S. (2008). The Stuff of Thought: Language as a Window into Human Nature. London: Penguin.

Prinz, J. J. (2004). Furnishing the mind: Concepts and their perceptual basis. Cambridge: MIT press.

Pulvermüller, F. (2013). How neurons make meaning: brain mechanisms for embodied and abstract-symbolic semantics. Trends in cognitive sciences, 17(9), 458–470.

Rabovsky, M., & McRae, K. (2014). Simulating the n400 erp component as semantic network error: Insights from a feature-based connectionist attractor model of word meaning. Cognition, 132(1), 68–89.

Řehŭřek, R., & Sojka, P. (2011). Gensim—statistical semantics in Python. Retrieved from genism.org.

Rosch, E. H. (1999). Principles of categorization. In E. Margolis, S Laurence (Eds.), Concepts: Core Readings, (pp. 189–206). Cambridge: MIT Press.

Rubinstein, D., Levi, E., Schwartz, R., & Rappoport, A. (2015). How well do distributional models capture different types of semantic knowledge? In Proceedings of the 53rd annual meeting of the association for computational linguistics and the 7th international joint conference on natural language processing (volume 2: Short papers) (Vol. 2, pp. 726–730).

Schölkopf, B., Herbrich, R., & Smola, A. J. (2001). A Generalized Representer Theorem. In D. Helmbold & B. Williamson (Eds.), Computational Learning Theory (pp. 416–426). Berlin, Heidelberg: Springer Berlin Heidelberg.

Simmons, W. K., Hamann, S. B., Harenski, C. L., Hu, X. P., & Barsalou, L. W. (2008). fMRI evidence for word association and situated simulation in conceptual processing. J Physiol Paris, 102(1-3), 106–119.

Sudre, G., Pomerleau, D., Palatucci, M., Wehbe, L., Fyshe, A., Salmelin, R., & Mitchell, T. (2012). Tracking neural coding of perceptual and semantic features of concrete nouns. Neuroimage, 62(1), 451–463.

Tissier, J., Gravier, C., & Habrard, A. (2018). Near-lossless Binarization of Word Embeddings. arXiv preprint arXiv:1803.09065.

Vo, M. L., Conrad, M., Kuchinke, L., Urton, K., Hofmann, M. J., & Jacobs, A. M. (2009). The berlin affective word list reloaded (bawl-r). Behavior Research Methods, 41(2), 534– 538.

Warglien, M., & Gärdenfors, P. (2013). Semantics, conceptual spaces, and the meeting of minds. Synthese, 190(12), 2165–2193.

Waskom, M., Botvinnik, O., O’Kane, D., Hobson, P., Ostblom, J., Lukauskas, S., & Qalieh, A. (2018). Mwaskom/seaborn: V0. 9.0 (july 2018). DOI: 10.5281/zenodo.592845.

Wehbe, L., Murphy, B., Talukdar, P., Fyshe, A., Ramdas, A., & Mitchell, T. (2014). Simultaneously uncovering the patterns of brain regions involved in different story reading subprocesses. PloS one, 9(11), e112575.

Xu, H., Murphy, B., & Fyshe, A. (2016, November). Brainbench: A brain-image test suite for distributional semantic models. In Proceedings of the 2016 Conference on Empirical Methods in Natural Language Processing (pp. 2017–2021).

